# Where species distribution models fail under occurrence-data contamination: calibration error concentrates at stream-network headwaters

**DOI:** 10.64898/2026.07.14.738364

**Authors:** Kristian Miok, Antonio V. Laza, Blaž Škrlj, Marko Robnik-Šikonja, Lucian Pârvulescu

## Abstract

Species distribution models (SDMs) increasingly inform conservation and biosecurity decisions in freshwater systems, where the reliability of its uncertainty estimates matters as much as its point predictions. Ensemble SDMs derive prediction intervals from across-replicate variance, but this variance captures systematic error only when replicates disagree about it, an assumption that fails when training data are contaminated with low-accuracy records, the norm in citizen-science datasets. Whether this failure is spatially uniform or concentrates in identifiable parts of a range is unknown. Using a panel of European freshwater crayfish spanning native headwater-associated species and invasive lowland colonizers, we show that contamination-induced calibration failure is strongly spatially structured: it concentrates at stream-network headwaters, the topological tops of the network, where upstream-aggregated predictors are structurally undefined, and scales with contamination severity, replicated across four species and both dominant ensemble protocols (replicate and consensus). The failure is driven by upward prediction bias, not by intervals failing to widen: contaminated ensembles overpredict suitability in headwaters, and because the bias is shared across ensemble members, the intervals do not flag it. This is a conservation-relevant blind spot, because headwaters are both refugia for threatened native crayfish and front lines for invasion; an SDM that silently overpredicts suitability there misdirects survey and management effort toward the segments where its predictions are least trustworthy. Standard leave-one-basin-out conformal calibration, the recommended panel-wide remedy, repairs marginal coverage but leaves headwaters undercovered, because a single calibration threshold is dominated by the abundant non-headwater segments. A group-conditional (Mondrian) variant, calibrating the two populations separately, restores reliable coverage in both at no extra cost and reallocates width where it is needed. We recommend network-position-stratified calibration as a default for ensemble SDMs in dendritic freshwater systems.

## 1. Introduction

Species distribution models (SDMs) are a central tool of quantitative biogeography and applied conservation, used to delineate protected areas, project range shifts, and prioritise surveillance for invasive species. In freshwater systems in particular, where field survey is costly and access uneven, managers increasingly lean on model predictions to decide where to look. The value of an SDM for these decisions rests not only on the accuracy of its point predictions but on the reliability of its uncertainty estimates: a 95% prediction interval is useful for decision-making only if it contains the truth close to 95% of the time. An interval that is narrow and confident-looking but systematically wrong is worse than no interval at all, because it invites misplaced trust.

The reliability of SDM predictions has been examined most thoroughly through the lens of spatial validation. Random cross-validation inflates apparent performance when observations are spatially autocorrelated, and a substantial literature now recommends spatially-blocked evaluation to obtain honest estimates of transferability (Roberts et al., 2017; Valavi et al., 2019; Ploton et al., 2020; Milà et al., 2022). That literature addresses one axis of unreliability, optimism from spatial autocorrelation between training and test points. It does not address a second axis that arises specifically from the way ensemble SDMs quantify uncertainty, nor whether the reliability of uncertainty estimates is itself spatially structured within a species’ range.

The de facto approach to uncertainty in applied SDM work is the ensemble: train a model many times under bootstrap resampling and varying random seeds, then derive prediction intervals from the empirical quantiles of the across-replicate distribution. The interval’s reliability is assumed to follow from the spread of the replicates, which holds only under a rarely tested condition, that across-replicate variance captures the systematic error of the modelling pipeline. When training data are contaminated with low-accuracy occurrence records, the norm rather than the exception in citizen-science-derived datasets such as GBIF (Beck et al., 2014; Zizka et al., 2019), the contamination injects a bias shared across all ensemble members, because they share the same training set. Internal disagreement between replicates does not reflect this shared bias, so the ensemble’s interval remains narrow even when its predictions are systematically wrong. On the crayfish panel analysed here, contamination drives the empirical coverage of nominally 95% ensemble intervals well below nominal across species, and split conformal prediction with leave-one-basin-out (LOBO) spatial folds restores panel-wide coverage to near-nominal.

That result is panel-level, and it leaves open the questions that matter most for how a manager should read a model surface. Where in space does the calibration failure concentrate? Is it uniform across a species’ range, or does it pool in ecologically identifiable subregions that could be flagged and treated with appropriate caution? And does the panel-wide conformal fix perform equivalently across those subregions, or does it repair the average while leaving a structured residual? A calibration scheme that achieves nominal coverage marginally but systematically misses in a defined sub-population is a coarser instrument than the conformal framework permits, and a refinement is available within the same framework (Vovk, 2003; Romano, Sesia & Candès, 2020).

Stream-network topology offers a natural and conservation-relevant axis along which to ask these questions. Freshwater SDMs operate over hydrographic networks in which environmental predictors are aggregated along the upstream catchment of each stream segment (Domisch et al., 2015; Amatulli et al., 2022). Headwater segments, segments at the topological top of the network, have no upstream catchment by definition, so upstream-aggregated predictors are structurally undefined for them, and the predictor space the model sees there is qualitatively different from the mid- and downstream network. Headwaters are also ecologically distinctive: they are strongholds and thermal refugia for threatened native crayfish and, at the same time, the front lines along which invasive crayfish and the crayfish plague they carry advance upstream. If contamination-driven bias concentrates where the predictor space is structurally distinct, headwaters are both the natural place to look and the place where getting the answer wrong carries the most direct management cost.

Concurrent work has begun to apply conformal prediction to SDMs (Poisot, 2025), and a broader literature on group-conditional and locally adaptive conformal prediction (Vovk, 2003; Romano et al., 2020; Tibshirani et al., 2019) provides the methodological scaffolding for stratified calibration. To our knowledge, no work has yet characterised the spatial structure of ensemble SDM miscalibration under realistic contamination, diagnosed the failure mode of standard split conformal in network-structured ecological systems, or evaluated topology-stratified alternatives in this setting.

Here we ask three questions on a panel of European freshwater crayfish. First, does ensemble SDM calibration failure under contamination concentrate at stream-network headwaters, and does it scale with contamination severity? Second, does standard LOBO conformal restore coverage uniformly across topological subregions, or leave a residual gap at network edges? Third, does group-stratified (Mondrian) calibration, with headwater status as the stratum, restore coverage simultaneously in both subregions, and at what efficiency cost? We answer all three on the four panel entities whose accessible ranges include substantial headwater representation, across both dominant ensemble protocols, and we verify that the pattern is not an artifact of feature-space sparsity. We find that contamination-driven calibration failure concentrates at headwaters, replicated in magnitude and direction across all four entities and both protocols; that standard LOBO conformal partially closes the gap but systematically undercovers headwaters; and that Mondrian LOBO restores nominal coverage in both populations while reallocating the width-inflation budget toward the population that needs it.

## 2. Methods

### 2.1 Study system and species panel

We tested whether ensemble SDM calibration fails systematically at the topological edges of stream networks using a panel of eight European freshwater crayfish entities. The panel spans native, alien, and historically translocated populations: *Astacus astacus*, *Austropotamobius torrentium* (pooled native populations), *Austropotamobius fulcisianus* (pooled), *Pacifastacus leniusculus* (alien North American invader), *Faxonius limosus* in both alien and native ranges, *Pontastacus leptodactylus* (pooled), and *Procambarus clarkii* in both alien and native ranges. We use the term *entity* throughout to denote a species-and-status combination treated as a distinct modelling unit. Occurrence records derive from a curated freshwater crayfish occurrence database; environmental predictors are drawn from the GeoTraits database (Petko et al. 2026) and combine local-scale (l_*) and upstream-aggregated (u_*) hydrographic, climatic, and substrate descriptors at the stream-segment scale, derived from the Hydrography90m and GeoFRESH platforms (Amatulli et al., 2022).

The four entities that carry the central analyses, *Astacus astacus*, *Austropotamobius torrentium*, *Faxonius limosus* (alien), and *Pacifastacus leniusculus* (alien), occur across stream networks with substantial proportions of headwater segments (35.5–44.7% of accessible-area pixels; total n = 84,035 pixels). This set spans the conservation contrast that motivates the paper: two native species of headwater and montane habitat, of conservation concern across much of their European range, and two North American invaders whose upstream spread threatens exactly those strongholds. They also encompass the topological transition we test, from headwater segments where upstream-aggregated covariates are structurally undefined to mid- and downstream segments where upstream context is available. The remaining four entities, two *Procambarus clarkii* populations, *Pontastacus leptodactylus*, and *Austropotamobius fulcisianus*, occupy networks with negligible headwater representation (0.3–1.0% of pixels each), reflecting habitat preference (*P. clarkii* is a lowland warm-water species; *P. leptodactylus* occupies large rivers and lakes) or the geographic structure of the accessible range. Because the headwater hypothesis cannot be tested where headwaters do not occur, we restrict the central analyses to the four headwater-bearing entities and report panel-wide statistics as a robustness check (Supporting information).

### 2.2 Surface generation and contamination protocol

We did not refit any model. The analyses operate on a fixed set of 30-replicate per-entity prediction surfaces, the deterministic benchmark surfaces, and the contamination protocol that produced them, described here in full. For each (entity, algorithm, predictor track, contamination level) cell, the pipeline generates 30 prediction surfaces by training the algorithm under bootstrap resampling with frozen hyperparameters and basin-stratified spatial cross-validation, then predicting onto every stream segment in the entity’s accessible area. Protocol A (replicate ensembles) uses two base algorithms, random forest and XGBoost, each treated as an independent 30-replicate ensemble. Protocol B (consensus ensembles) uses four algorithms, a generalised linear model, a generalised additive model, random forest, and XGBoost, each fit once, with the 95% interval taken from the quantiles of the four predictions, following standard ensemble-SDM practice (Araújo & New, 2007); its calibration reference is the consensus of the four clean-data predictions. The contamination protocol injects low-accuracy occurrence records, flagged as imprecise in the master database, at 3%, 10%, and 20% of the training set. A deterministic benchmark surface is produced for each (entity, algorithm) pair under uncontaminated training data and serves as the calibration reference.

The present study uses the combined predictor track (398 candidate features after track-specific subsetting; 291–299 features after the missingness and correlation filter applied by the pipeline). We analyse the combined track only, because it is the sole track on which the headwater partition is meaningful: under upstream-only predictors, headwater segments have almost no defined predictor at all, so any effect would be trivially driven by absent inputs rather than shared bias; under local-only predictors the structural headwater/non-headwater distinction does not arise. Combined is also the full predictor set practitioners use by default. We restrict the central analyses to the low-accuracy contamination axis and report all three contamination levels (3%, 10%, 20%) to characterise the contamination–response relationship, focusing on the most severe level (20%) for the mechanistic and correction analyses. Per-replicate prediction surfaces, the predictor matrix used for model fitting, and the deterministic benchmark surface for each cell were extracted from the pipeline and saved as pixel-aligned arrays. All extractions and analyses were performed without retraining; the predictor matrix in our analyses is byte-identical to the matrix the model was trained on.

### 2.3 Headwater identification

We define a stream segment as a headwater if it has no resolved upstream catchment in the hydrographic network used for predictor aggregation. Operationally, a segment is classified as headwater if at least 20 of its upstream-aggregated (u_*) features are missing in the predictor matrix, against a panel norm in which non-headwater segments have zero or near-zero missingness across upstream features. The classification is sharp: across the four entities in the headwater panel, 97.8–98.1% of headwater-classified rows have only upstream features missing, while 1.6–2.6% additionally have one or more local features missing (interpreted as concurrent local data-quality issues at headwater locations rather than a separate population). The bimodality is essentially complete: segments are either fully observed (no missing features, 55.3–57.0% of pixels) or heavily missing (≥ 20 missing upstream features, 35.5–44.7% of pixels); fewer than 0.1% of pixels fall between these categories.

The structural-missingness definition aligns with the topological definition of a headwater (a segment with no upstream catchment) up to the bimodality reported above, and we use the two terms interchangeably. We do not impose an additional spatial filter (e.g., Strahler order) because the structural-missingness signal already isolates the population of interest cleanly; we note that in predictor pipelines that do not use upstream aggregation, an explicit topological definition would be required instead. For analyses requiring complete predictor matrices, we impute missing cells with the column median over the full entity panel; this preserves all rows and columns and tracks which rows were imputed.

### 2.4 Per-pixel calibration error

For each (entity, algorithm, contamination level) cell, we compute per-pixel ensemble prediction intervals as the 2.5% and 97.5% empirical quantiles across the 30 contaminated replicates. Empirical coverage at each pixel is the binary indicator that the deterministic benchmark prediction lies inside the interval. We report two pixel-level error measures. The miscalibration indicator is the binary 1[benchmark ∉ interval] and aggregates by averaging to the empirical coverage gap from nominal. The miscalibration distance is the signed distance from the benchmark to the nearest interval edge (zero if inside; positive if outside) and captures the magnitude of the overrun. Both measures are computed against the deterministic benchmark, the prediction the same model would produce under uncontaminated training, not against held-out occurrence labels; the question we ask is whether the contaminated ensemble’s interval contains the prediction the clean model would have made. This framing isolates contamination-induced calibration error from baseline model error.

We further decompose the per-pixel error into bias and width components. Per-pixel bias is the difference between the across-replicate mean prediction and the benchmark; per-pixel interval width is the difference between the 97.5% and 2.5% quantiles. Comparing bias and width between headwater and non-headwater populations isolates whether the calibration failure arises from prediction shift (bias) or from intervals failing to widen (variance underestimation).

### 2.5 Conformal calibration

We apply two variants of split conformal calibration with leave-one-basin-out (LOBO) folds. The non-conformity score for a pixel is the distance from the benchmark to the nearest interval edge, identical to the miscalibration distance defined in §2.4. Under split conformal prediction, the calibrated interval at a test pixel is widened on both sides by the (1 − α) finite-sample quantile of the non-conformity scores on the calibration set, with α = 0.05 throughout. Under LOBO cross-validation, the calibration set for each test basin consists of all pixels in the remaining basins; the calibrated quantile q^ is computed once per fold and applied to all test pixels in that fold. The use of basin-level folds means it already controls for spatial autocorrelation in the sense of Roberts et al. (2017).

We evaluate two stratification schemes within this framework. Standard LOBO conformal computes one calibration quantile per fold, pooled across all calibration pixels, and applies it uniformly to all test pixels. Mondrian LOBO conformal (Vovk, 2003; Romano et al., 2020) computes two calibration quantiles per fold, one from headwater calibration pixels, one from non-headwater calibration pixels, and applies them respectively to headwater and non-headwater test pixels. The two schemes differ by a single line of implementation and have identical computational cost. Per-fold quantiles use the finite-sample correction of Lei et al. (2018), specifically the ⌈(n+1)(1−α)⌉/n-th order statistic of the non-conformity scores. After each fold we record the corrected per-pixel intervals, recompute coverage and width on the held-out basin, and aggregate across folds, reporting empirical coverage and median interval width separately for headwater and non-headwater pixels, together with the width-inflation factor (post- over pre-correction median width within each population).

### 2.6 Robustness: controlling for feature-space sparsity

A natural alternative explanation for elevated calibration error at headwaters is sparsity: headwater segments may sit in undersampled or extreme regions of the predictor space, where any model performs poorly for reasons unrelated to network topology. To test this, we computed a local density estimate for each pixel from its k = 50 nearest neighbours in the standardised, median-imputed predictor matrix (the inverse of mean k-nearest-neighbour distance), and asked whether the headwater–non-headwater calibration gap persists after controlling for it. We report the gap within matched local-density strata and the coefficient of headwater status in a logistic regression of per-pixel miscalibration on headwater status and local density.

As a secondary, exploratory analysis we additionally examined whether the local intrinsic dimension of the predictor space, a geometric descriptor of how many effective dimensions the data occupy locally, estimated with the TwoNN method (Facco et al., 2017), provides further within-headwater stratification of calibration risk. This analysis is reported in the Supporting information and is not part of the main results; we summarise it briefly in the Discussion.

### 2.7 Software and reproducibility

All analyses were implemented in Python 3.12. All analysis code, conformal calibration, topology-aware extensions, panel-level diagnostics, robustness analyses, and figure generation, is released in the accompanying repository (see the Data availability statement below). The repository includes the full analysis pipeline and a synthetic-data smoke test that verifies the calibration pipeline end-to-end on a constructed dataset with known ground-truth structure. All numerical results are reproducible from the released code on the archived panel surfaces. Analyses ran on a 2024 Apple M3 MacBook in approximately 45 minutes of wall time.

## 3. Results

### 3.1 Calibration failure concentrates at headwaters and scales with contamination

Across the four entities with substantial headwater populations, contamination of training data induced a coverage gap between headwater and non-headwater stream segments that scaled monotonically with contamination severity (Table 1). On random forest, mean headwater coverage fell from 0.871 at 3% contamination to 0.621 at 20%, while non-headwater coverage fell only from 0.895 to 0.779 over the same range; the mean coverage gap (non-headwater minus headwater) grew from +0.024 at 3% to +0.158 at 20%. The pattern was sharper on XGBoost: at 20% contamination, mean headwater coverage was 0.467 against non-headwater 0.663, a gap of +0.196, with all four entities showing a positive headwater–non-headwater gap and the gap increasing monotonically with contamination level in every entity.

**Table 1.** Empirical coverage of contaminated-ensemble 95% prediction intervals by population, ensemble protocol, and contamination level. Random Forest and XGBoost rows are Protocol A (replicate ensembles); Consensus rows are Protocol B (four-algorithm consensus). Coverage is the empirical proportion of stream segments at which the deterministic benchmark prediction lies inside the contaminated-ensemble interval. Means across the four entities in the headwater panel (n = 84,035 segments).

| Algorithm | Level | Headwater coverage | Non-headwater coverage | Gap (NH – HW) |
| --- | --- | --- | --- | --- |
| Random Forest | L3 | 0.871 | 0.895 | +0.024 |
| Random Forest | L10 | 0.736 | 0.850 | +0.114 |
| Random Forest | L20 | 0.621 | 0.779 | +0.158 |
| XGBoost | L3 | 0.717 | 0.836 | +0.119 |
| XGBoost | L10 | 0.551 | 0.736 | +0.185 |
| XGBoost | L20 | 0.467 | 0.663 | +0.196 |
| Consensus (4-algo) | L3 | 0.954 | 0.987 | +0.033 |
| Consensus (4-algo) | L10 | 0.820 | 0.940 | +0.120 |
| Consensus (4-algo) | L20 | 0.706 | 0.884 | +0.178 |

The mechanism was upward prediction bias rather than narrowed ensemble variance (Table 2). At 20% contamination on random forest, the mean bias of the contaminated ensemble (ensemble mean minus benchmark) was +0.106 in headwater segments and +0.032 in non-headwaters, a gap of +0.074 in the same direction as the coverage gap. Median interval widths were similar between populations (0.298 vs. 0.244 on random forest at 20%; 0.345 vs. 0.272 on XGBoost), and headwater intervals were modestly wider, not narrower, than non-headwater intervals. Ensemble variance therefore did widen with contamination, but did so roughly equally in both populations and not enough to compensate for the headwater-concentrated bias. The four entities outside the headwater panel had headwater proportions below 1% and contributed essentially no signal to the comparison; we treat them as not testing the hypothesis rather than as negative results (Supporting information).

**Table 2.** Bias and width decomposition at 20% contamination. Bias is the mean of (ensemble mean − benchmark) within each population; width is the median per-pixel 97.5%–2.5% interval.

| Entity | Algo | Bias HW | Bias non-HW | Width HW | Width non-HW |
| --- | --- | --- | --- | --- | --- |
| Faxonius limosus (alien) | RF | +0.091 | +0.011 | 0.291 | 0.281 |
| Austropotamobius torrentium | RF | +0.110 | +0.046 | 0.234 | 0.158 |
| Astacus astacus | RF | +0.130 | +0.047 | 0.298 | 0.259 |

| Entity | Algo | Bias<br>HW | Bias non-HW | Width<br>HW | Width non-HW |
| --- | --- | --- | --- | --- | --- |
| Pacifastacus leniusculus<br>(alien) | RF | +0.093 | +0.021 | 0.190 | 0.097 |
| Faxonius limosus (alien) | XGB | +0.144 | +0.045 | 0.345 | 0.272 |
| Austropotamobius torrentium | XGB | +0.166 | +0.063 | 0.258 | 0.169 |
| Astacus astacus | XGB | +0.183 | +0.077 | 0.319 | 0.255 |
| Pacifastacus leniusculus<br>(alien) | XGB | +0.131 | +0.038 | 0.215 | 0.122 |

### 3.2 The headwater effect is not explained by feature-space sparsity

The concentration of calibration failure at headwaters is not a sparsity artifact. Within matched local-density strata (feature-space density quintiles), the headwater–non-headwater miscalibration gap persisted in the great majority of strata and was only modestly attenuated relative to the unstratified gap: the grand-mean density-matched gap across the four entities was +0.146, against unstratified gaps of +0.080 to +0.213. In a logistic regression of per-pixel miscalibration on headwater status and local density, the coefficient of headwater status was positive in all four entities (mean +0.705), whereas the coefficient of local density was small and inconsistent in sign across entities (+0.05, +0.36, +0.21, −0.27). Local density therefore neither explains away the headwater effect nor exerts a consistent effect of its own. The elevated calibration error at network edges reflects the structural distinctness of the headwater predictor space, where upstream-aggregated covariates are undefined, rather than merely lower sampling density there. Per-stratum values are reported in the Supporting information.

### 3.3 Standard LOBO conformal undercovers headwaters and misallocates width

Standard LOBO conformal calibration restored non-headwater coverage to nominal across all four entities and both algorithms but systematically undercovered headwaters (Figure 1). On random forest, mean post-LOBO non-headwater coverage at 20% was 0.965, at or above nominal in every entity, while mean post-LOBO headwater coverage was 0.887, with all four entities below nominal (0.857–0.923). On XGBoost the gap was sharper still: 0.969 non-headwater versus 0.892 headwater. Standard split conformal therefore restored the marginal coverage of the panel as a whole while leaving a systematic conditional miscoverage at the topological edges of the network.

**Figure 1.**
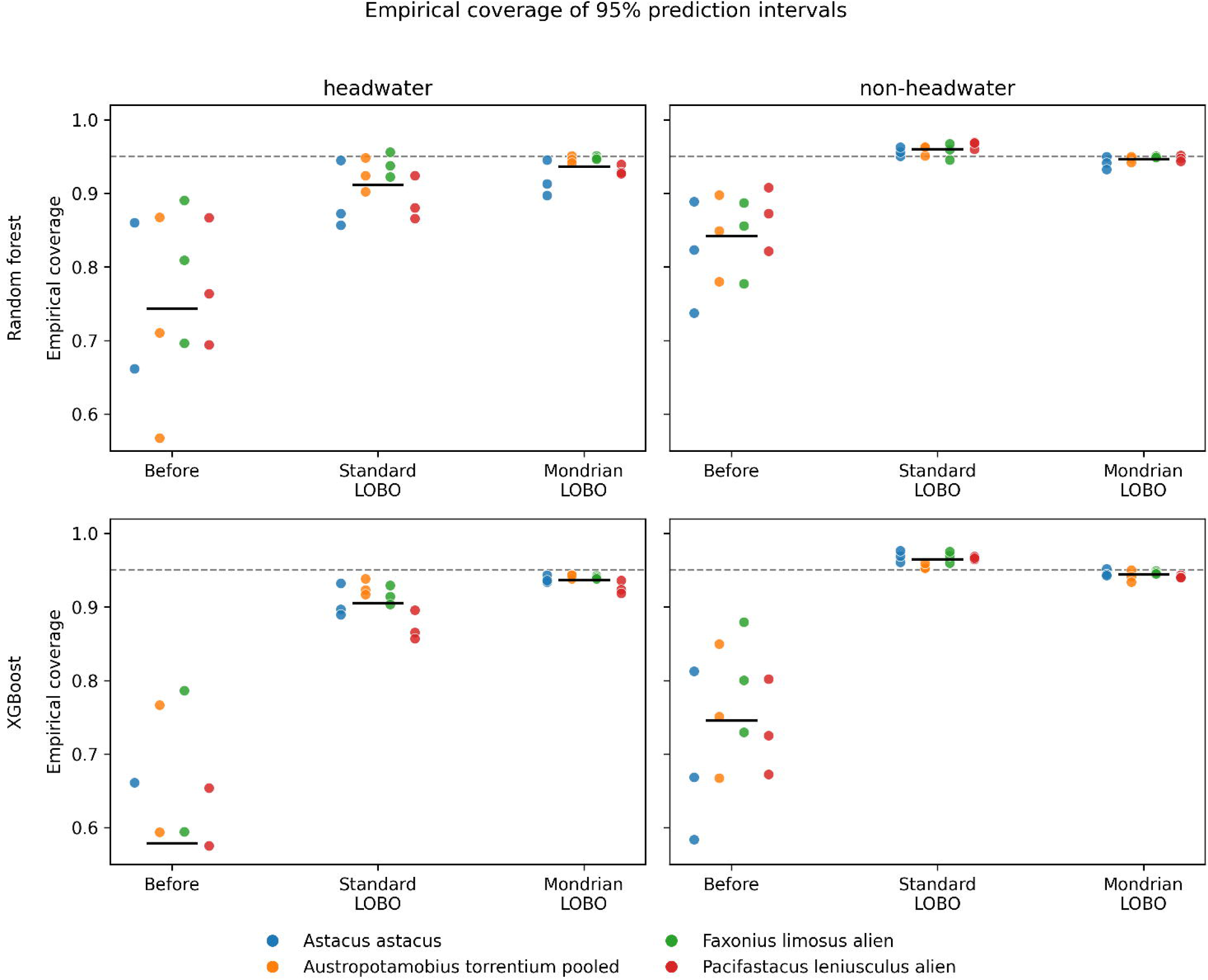
Empirical coverage of 95% prediction intervals before correction, after standard LOBO split conformal, and after Mondrian LOBO split conformal, separately for headwater and non-headwater stream segments. Top row: random forest; bottom row: XGBoost. Each point is one (entity, contamination level) combination; horizontal dashed line at 0.95 marks nominal coverage. Standard LOBO restores non-headwater coverage to nominal but undercovers headwaters; Mondrian LOBO restores both populations simultaneously.

**Figure 2.**
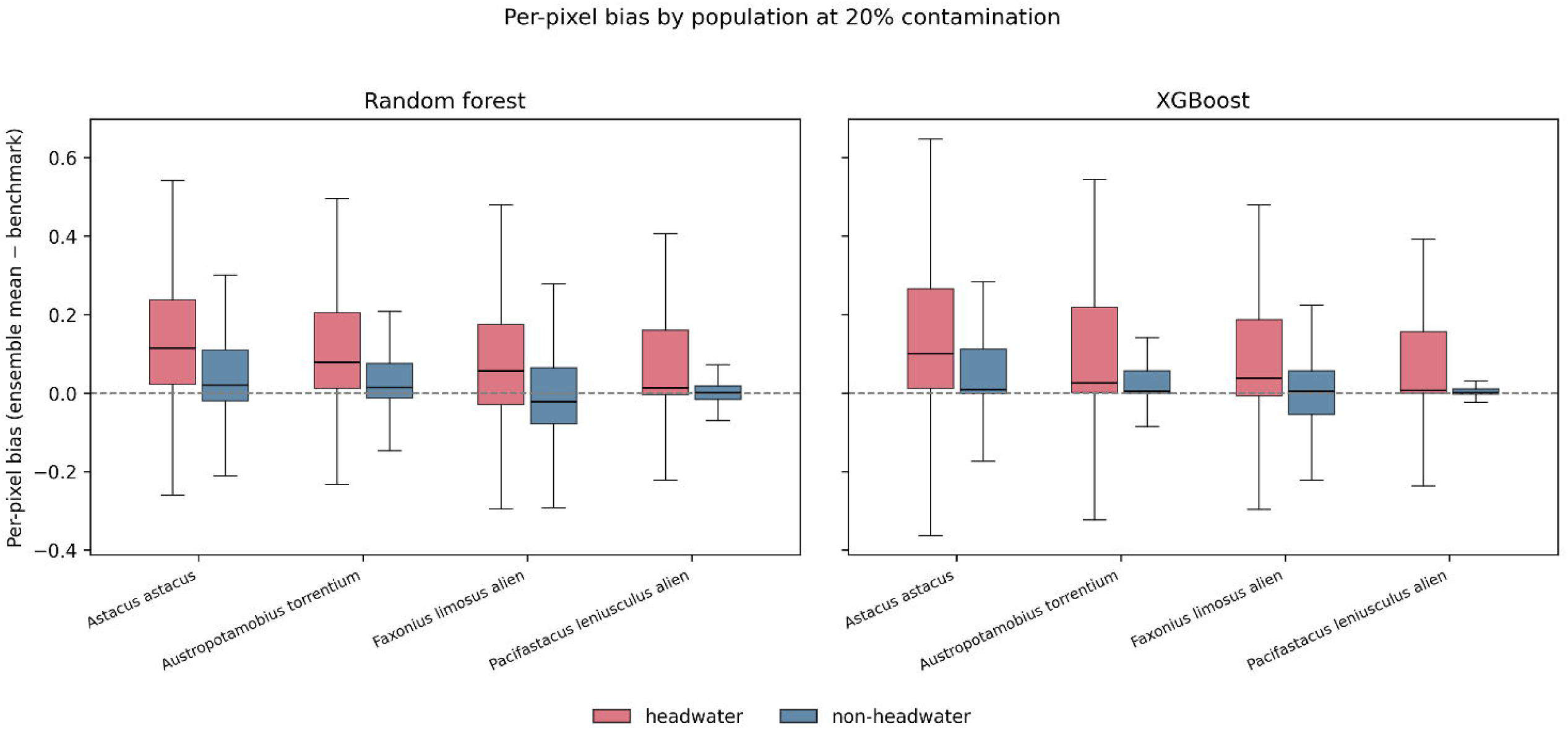
Per-pixel bias (ensemble mean − benchmark) by population at 20% contamination, four entities × two algorithms. Boxplots show the distribution of per-pixel bias within headwater and non-headwater segments. Headwater segments show consistently larger upward bias across all eight cells.

**Figure 3.**
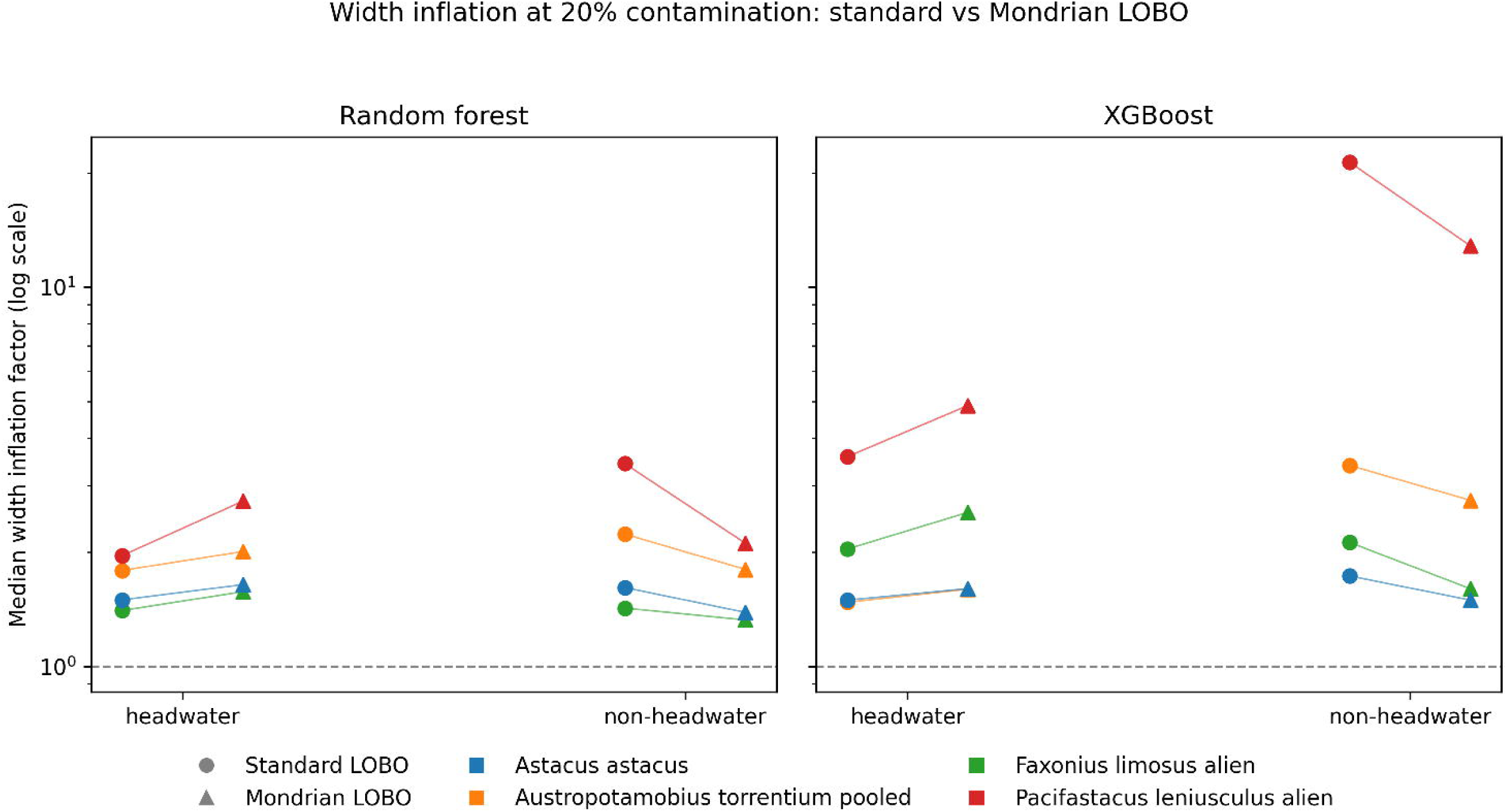
Median width inflation factor under standard LOBO and Mondrian LOBO conformal calibration, by population, at 20% contamination. Mondrian increases inflation in headwaters (where it is needed to close the coverage gap) and decreases it in non-headwaters (where standard LOBO over-corrected to the pooled tail). Net total width budget is lower under Mondrian in 7 of 8 entity–algorithm cells.

Width inflation under standard LOBO was allocated inversely to where it was needed. In three of four entities at 20% contamination on random forest, non-headwater intervals were inflated more than headwater intervals (e.g., *Pacifastacus leniusculus* alien: 1.96× headwater vs. 3.42× non-headwater); the same pattern held under XGBoost, where the contrast was more extreme (*Pacifastacus* alien: 3.57× headwater vs. 21.32× non-headwater). The pooled non-conformity quantile is dominated by the larger non-headwater calibration set, whose errors are on average smaller in magnitude, so the resulting q^ is tuned to the easier population: headwater pixels, with larger errors but a smaller share of the calibration set, are under-corrected, while non-headwater pixels are over-corrected to match the pooled tail.

### 3.4 Mondrian LOBO conformal restores coverage and reallocates width

Group-stratified (Mondrian) LOBO conformal calibration restored headwater coverage to within 1.6 percentage points of nominal at 20% contamination on random forest (mean 0.932; all four entities 0.913–0.947) and within 1.6 points on XGBoost (mean 0.934; range 0.919–0.943), while simultaneously maintaining non-headwater coverage at near-nominal levels (mean 0.942 on random forest, 0.941 on XGBoost). The non-headwater quantile under Mondrian is very close to the pooled quantile under standard LOBO, which is why marginal coverage in the easier population is preserved; the Mondrian fix closes the conditional miscoverage gap without sacrificing it.

The width-budget reallocation was substantial (Supporting information). Under Mondrian, headwater intervals received larger inflation than under standard LOBO in all four entities under both algorithms (e.g., Pacifastacus alien on XGBoost: 3.57× → 4.87×), while non-headwater intervals received smaller inflation (21.32× → 12.85×). The net effect across the panel was a 23–62% reduction in non-headwater width inflation, paid for by a 12–37% increase in headwater width inflation, and the total width budget, the summed inflated width over all pixels, was lower under Mondrian than under standard LOBO in all four entities under XGBoost and in three of four under random forest. Mondrian calibration therefore does not merely trade coverage between populations; on this panel it delivers reliable coverage in both while producing narrower intervals overall in most cells.

The spatial structure of this correction is shown for one representative drainage basin in Figure 4. Before correction, uncovered segments cluster on the headwater tips of the network in both species. Standard LOBO clears them almost entirely for *Austropotamobius torrentium* but leaves a visible headwater residual for *Pacifastacus leniusculus*; Mondrian calibration reduces that residual further. The concentration documented panel-wide above is therefore legible as a property of network topology in a single basin.

**Figure 4.**
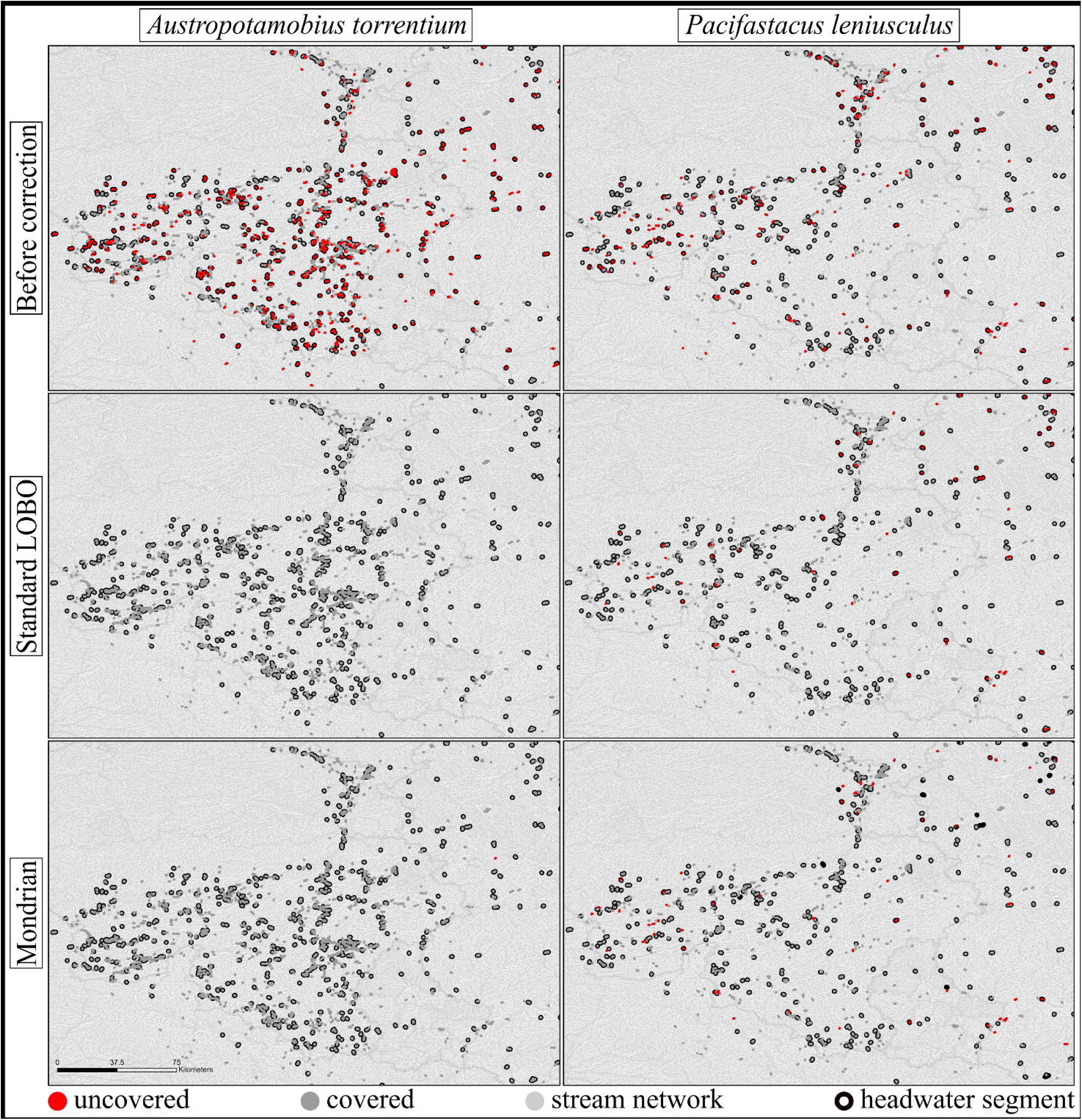
Spatial concentration of prediction-interval miscoverage at stream-network headwaters, shown for one representative drainage basin (Hydrography90m basin 1173482) at 20% contamination (random forest). Columns: *Austropotamobius torrentium* (native montane refugium species) and *Pacifastacus leniusculus* (invader). Rows: the contaminated ensemble before correction, after standard leave-one-basin-out (LOBO) split conformal calibration, and after group-conditional (Mondrian) LOBO calibration. Red segments are uncovered by that regime’s 95% prediction interval; grey segments are covered. Headwater segments are outlined with a black halo. The full physical stream network is shown in light grey beneath; only segments that entered the SDM (above the stream-order and flow-accumulation threshold) are coloured, which is why the modelled network appears discontinuous. Headwater coverage in this basin runs 0.40 → 0.87 → 0.93 for *A. torrentium* and 0.39 → 0.58 → 0.77 for *P. leniusculus* across the three regimes (non-headwater: 0.71 → 0.94 → 0.91 and 0.68 → 0.82 → 0.77). This is one representative basin; *P. leniusculus* headwaters are not fully corrected here (0.77), and panel-wide coverage across all basins is reported in the main text.

### 3.5 Both findings replicate under the four-algorithm consensus ensemble

All results above use Protocol A. To test whether the pattern is specific to that construction, we repeated the full analysis under Protocol B, the four-algorithm consensus ensemble. The headwater concentration, its contamination scaling, and the LOBO-versus-Mondrian contrast all replicate. Averaged across the four entities, pre-correction headwater coverage fell from 0.954 at 3% contamination to 0.706 at 20%, against non-headwater coverage of 0.987 and 0.884, the same headwater-concentrated gap seen under Protocol A. At 20% contamination, standard LOBO again undercovered headwaters (mean 0.876, below nominal in all four entities) while restoring non-headwater coverage (0.969), and Mondrian LOBO again closed the headwater gap (0.931) while holding non-headwater coverage near nominal (0.936). That the failure survives the structural diversity of four model families, not only bootstrap resampling of one, supports the proposed mechanism: at headwaters every algorithm lacks upstream context and shares the same upward projection, so even a consensus does not disagree about the error. Because the consensus interval rests on only four predictions, its pre-correction non-headwater coverage is mildly conservative (0.94–0.99), and Mondrian brings it closer to nominal.

## 4. Discussion

Three findings stand out. First, where ensemble SDMs fail under realistic contamination of training data, they fail systematically at the topological edges of the stream network rather than uniformly across the prediction surface. The headwater concentration is replicated across four ecologically heterogeneous entities, two native European species associated with headwater and montane habitat, two North American invaders, under both ensemble protocols, replicate ensembles of two base algorithms and a four-algorithm consensus, at all three contamination levels, and scales monotonically with contamination severity. Second, the failure mode is upward prediction bias rather than narrowed ensemble variance: contaminated ensembles overpredict suitability in headwaters, and because that bias is shared across ensemble members, bootstrap replicates and distinct algorithms alike, neither across-replicate nor across-algorithm disagreement sees it, and the intervals stay narrow. Third, the standard split conformal correction proposed for the panel-wide problem restores marginal coverage but leaves a systematic conditional miscoverage at network edges; a small, existing modification of the conformal framework, Mondrian stratification by headwater status, closes that gap and, on this panel, simultaneously narrows intervals overall.

The mechanism is straightforward and, we suspect, general to network-aggregated predictor pipelines. Headwater segments are the region of predictor space where upstream-aggregated covariates are structurally undefined, forcing the model to predict from local-scale features alone. Contamination introduces low-accuracy occurrence records that are systematically more likely to fall in accessible, lower-elevation, mid-to-downstream habitat, where reporting is easier. The model fits this contamination-shifted distribution and, lacking upstream context, projects lower-elevation habitat associations onto headwater segments where they do not apply. The bias is shared across replicates because all replicates see the contamination; the variance is small because all replicates make the same shared mistake. This is the canonical mechanism for over-confident ensemble miscalibration in the conformal-prediction literature (Angelopoulos & Bates, 2023), and it is precisely why post-hoc calibration against held-out data, rather than internal ensemble disagreement, can repair it.

Standard LOBO conformal repairs the marginal failure but inherits the structural blind spot, because a single pooled non-conformity quantile is dominated by the abundant, lower-error non-headwater segments. This is a direct consequence of conformal theory: split conformal guarantees marginal coverage but not group-conditional coverage when the calibration distribution differs across groups, and Vovk (2003) introduced Mondrian conformal precisely to restore conditional guarantees. Our use of Mondrian conformal here is methodologically modest, the framework is over twenty years old, but is, to our knowledge, the first time it has been motivated by an empirically demonstrated conditional miscoverage on real ecological data and shown to close the gap without sacrificing marginal coverage. The contribution of the present paper is the empirical and applied step from “there is a panel-wide calibration problem” to “the problem has a specific, conservation-relevant spatial structure that an existing tool repairs cleanly.” We also confirmed, through density-matched comparison and density-controlled regression, that the headwater concentration is not an artifact of feature-space sparsity; it tracks the structural distinctness of the headwater predictor space.

As a secondary and exploratory line, we examined whether local intrinsic dimension of the predictor space provides further stratification of calibration risk *within* the headwater population. We found a modest, replicated positive association (reported in the Supporting information), consistent with the interpretation that headwater segments in geometrically more complex regions of the predictor space lie further from the cases the model was effectively trained on. The signal is real but small, and we report it as supportive of the broader claim that calibration risk is spatially and structurally patterned rather than uniform, not as a stand-alone diagnostic.

Why this blind spot matters is inseparable from the ecology of the taxa it affects. For the two native species in our panel, the noble crayfish *Astacus astacus* and the stone crayfish *Austropotamobius torrentium*, headwaters are increasingly the habitat of last resort. *A. astacus* is listed as Vulnerable on the IUCN Red List and on Annex V of the EU Habitats Directive; *A. torrentium*, globally Data Deficient but in documented decline, is listed on Annexes II and V. Both are highly susceptible to crayfish plague (*Aphanomyces astaci*), carried asymptomatically by the two invaders in the same panel, *Pacifastacus leniusculus* and *Faxonius limosus*. As these plague-carrying invaders spread through accessible lowland and mid-order reaches, native populations contracted upstream, persisting in cold, isolated headwaters where low temperatures and dispersal barriers slow both invader and pathogen. The same cold-water isolation that makes *A. torrentium* an obligate montane specialist makes headwaters the thermal refugia into which both natives are pushed as lowland climates warm. For these species, the stronghold and the network edge have become the same place.

This ecology gives the statistical result its bite, and it does so asymmetrically, between the two native species and the two invaders. The failure we document is a single kind of error, an overconfident overprediction of suitability at headwaters, but its management consequences run in opposite directions. For the native refugium species, a model that confidently overstates headwater suitability paints the last strongholds as more secure than they are, directing monitoring away from precisely the reaches where a plague outbreak or a failed year-class would go undetected; the blind spot falls on the populations of highest conservation priority. For the invaders, the same directional error inflates apparent invasibility at the upstream front, the reaches where containment decisions of barriers, surveillance, and rapid response are made, while the narrow interval hides how little the model knows there. In both cases the danger is not that the point prediction is wrong, which is unavoidable, but that the reported uncertainty says it is right.

Standard LOBO conformal calibration does not remove this danger: it closes the marginal gap while leaving a residual headwater-specific miscoverage of several percentage points at moderate-to-severe contamination, a gap that falls, again, on exactly the segments where the ecological stakes are highest. The headwater-stratified Mondrian variant is the same pipeline with the calibration step computed twice, and it restores honest intervals where they matter most. For SDMs operating over hydrographic networks with upstream-aggregated predictors, and especially where those models inform crayfish conservation or invasion-risk assessment, we recommend it as a default.

Several limitations bound these conclusions. The panel is four headwater-bearing crayfish entities in European stream networks; whether the concentration generalises to other freshwater taxa with comparable network structure, or to terrestrial systems with analogous topological edges, is open. The mechanism we propose is general, but its empirical magnitude is system-specific. The contamination model we test is a single low-accuracy axis; other mechanisms (positional jitter, identification error, sampling bias) may produce different spatial structures of failure. The headwater partition is defined by the structural-missingness signature of upstream-aggregated features, so elsewhere an explicit topological definition (e.g., Strahler order 1) would be needed. Finally, we evaluate the spatial-fold split-conformal scheme; full conformal or jackknife+ would give tighter guarantees at higher cost (Barber et al., 2021). Nor do we claim Mondrian is right where the relevant axis of conditional miscoverage is continuous rather than categorical; there, conformalised quantile regression (Romano et al., 2020) may be preferable. We chose Mondrian because the headwater partition is sharp, its interpretation is clean, and its cost is one line of code.

Standard split conformal calibration restores panel-wide coverage but, as we have shown, leaves a structured residual at the topological edges of the network that a group-conditional refinement removes. The trustworthy quantification of SDM uncertainty is, in our view, a research programme rather than a single contribution, and we hope this paper marks one step within it.

## Acknowledgements

Acknowledgements and funding statements are provided on the separate title page and are omitted here to preserve double-blind review.

## Data availability

All analysis code and the pixel-aligned prediction surfaces required to reproduce the results, including the Protocol B robustness analysis, are available for peer review at https://zenodo.org/records/21130430?preview=;1&token=;eyJhbGciOiJIUzUxMiJ9.eyJpZCI6IjRhZTMxZTE4LWI4NWUtNDY4My05ZWQ0LTM0N2E4OTA0YTMwZCIsImRhdGEiOnt9LCJyYW5kb20iOiI3Yjc4NDYwNTQ2ZWZmNmI2ZGE0N2ZhYWI2ZWY0YjhkZiJ9.7DEm-er5UDfzb7Z2NuyPMMFtc1pPElPs5fNFCJOL4iFNJWauPwtiuq8u87uQhN7PA788qRCRu3J723ae1BBg8g (anonymised review link). A citable release with a permanent identifier will be minted upon acceptance.

